# Genealogies under purifying selection

**DOI:** 10.1101/2024.10.15.618444

**Authors:** Ksenia A. Khudiakova, Florin Boenkost, Julie Tourniaire

## Abstract

Selection against deleterious mutations, called purifying selection, plays a central role in evolution and acts in all populations. It is known that the genetic patterns observed in genomic regions undergoing purifying selection differ from those resulting from neutral evolution. However, a comprehensive understanding of the underlying mechanisms shaping those patterns is still lacking.

In the present work, we use simulations combined with a genealogical approach to identify the effect of purifying selection on the ancestry and thus on the genetic diversity. Our analysis relies on the postulate that the genealogy belongs to the universality class of Beta-coalescents. Under this assumption, we derive statistics measuring the distortion of the genealogy. This approach allows us to consider a wide range of regimes (i.e. arbitrary selection and mutation strengths) and uncover a rich phase diagram. We find that, for strong selection, the limiting genealogy is given by Kingman’s coalescent on a polynomial timescale. As selection gets weaker, Muller’s ratchet starts operating, setting off the emergence of multiple mergers in the genealogical structures. Our results show that while multiple-merger coalescents are often interpreted as the signature of selective sweeps in rapidly adapting populations, these structures can also appear in the context of Muller’s ratchet.

## 1 Introduction

Deleterious mutations occur in every population, in every generation. Not only do they directly impact the population’s fitness distribution, but they also have indirect effects on the surrounding loci. These indirect effects translate into a local reduction in genetic diversity: neutral variation is purged together with deleterious mutations by natural selection, a phenomenon called background selection [Charlesworth et al., 1993]. This diversity loss becomes more drastic as the rate of recombination decreases so that non-recombining parts of the genome – such as Y-chromosome and mitochondrial DNA – are the most affected ones. In that sense, deleterious mutations constitute a key factor in evolution. Qualitatively, this phenomenon is similar to another form of linked selection, known as genetic hitchhiking, where neutral variants surrounding a beneficial mutation rapidly reach fixation [Maynard Smith and Haigh, 1974].

From a backward-in-time perspective, diversity loss is usually related to distortions in genealogical structures. Under neutrality, the genealogy of a population of size *N* is described by Kingman’s coalescent [Kingman, 1982] on a timescale of order *N*. This coalescent process involves only binary mergers, resulting in a relatively high genetic diversity. In contrast, parts of the genome under directional selection exhibit lower genetic diversity [Maynard Smith and Haigh, 1974, Charlesworth et al., 1993]. In this respect, Kingman’s coalescent constitutes a null-model for the detection of deviations from neutrality. However, relying on the reduction in genetic diversity alone cannot distinguish between negative and positive selection. Therefore, understanding the structure of genealogies under different forms of selection is crucial for differentiating the two patterns of reduced genetic diversity.

The genealogies in asexual populations under continual adaptation were extensively studied in models of fitness waves [Brunet et al., 2007, Neher and Hallatschek, 2012, Brunet and Derrida, 2012, Schweinsberg, 2017].

In this framework, selection induces a high reproductive variance, which, in turn, gives rise to genealogies with multiple mergers [Schweinsberg, 2003]. These multiple-merger genealogies stand in sharp contrast with binary trees obtained in the neutral case.

The effects of purifying selection on genealogies are less well understood. Assuming full linkage and equal selection coefficients at all loci, the population’s fitness distribution is well-approximated by a Poisson distribution with parameter *µ/* |*s*|, where *µ* is the genome-wide deleterious mutation rate and *s* is the selection coefficient [Haigh, 1978]. Under strong selection, the fitness distribution maintains a stable shape. When selection is weak, the fitness class with zero mutation can be lost due to stochastic effects. In asexual populations, this loss is irreversible. We say that the ratchet clicks. After the click, the fittest individuals are those carrying a single deleterious mutation, resulting in a shift of the fitness distribution to lower values. This dynamics, known as Muller’s ratchet, can continue until the mean fitness declines to a point where the population goes extinct [Muller, 1964, Haigh, 1978]. It is reasonable to believe that stationary and non-stationary scenarios give rise to different genetic patterns and genealogies. To the best of our knowledge, the genealogies in these two cases were treated separately so far.

In the stationary scenario, genealogies were studied using the so-called *fitness-class coalescent* approach. This method was developed to explain the reduction in genetic diversity observed under strong purifying selection [Nicolaisen and Desai, 2012, Walczak et al., 2012]. In this framework, ancestral lineages are traced back through the fitness distribution. In each class, a lineage can either move to a fitness class with a lower number of mutations, or coalesce with another lineage located in the same class. Using this model, several statistics of genetic diversity, including coalescence probabilities and pairwise heterozygosity, were derived [Nicolaisen and Desai, 2012, Walczak et al., 2012]. This approach shows that, most of the time, the lineages travel back to the fittest class before coalescing in this class. As a consequence, the resulting statistics of genetic diversity resemble that of a neutral model with reduced effective population size corresponding to the size of the fittest class. In the present work, we achieve similar results in the strong selection regime using coalescent theory. Recently, [Strütt et al., 2024] extended the *fitness-class coalescent* approach to the regime of weak selection. This adjustment requires to modify the lineage dynamics: when the ratchet operates, lineages are allowed to travel to fitness classes containing more mutations. These events correspond to a shift in the fitness distribution resulting from a click of the ratchet. Using this framework, the authors compute the distribution of pairwise coalescent times and the inferred population size.

Here, we use a numerical approach based on standard coalescent theory to characterise the genealogy of a population undergoing purifying selection, in both strong and weak selection regimes. This approach allows us to overcome the need to treat stationary and non-stationary regimes separately. Moreover, it numerically captures the emergence of multiple mergers as selection gets weaker. We apply this method to the Wright-Fisher model with selection and mutation. In the strong selection regime, our results are in line with the reduced “effective population size” prediction [Charlesworth et al., 1993], suggesting that the genealogy in the strong selection regime is described by Kingman’s coalescent with a reduced population size. Moreover, they shed light on the weak selection regime, where we observe multiple-merger coalescents, akin to those found in rapidly adapting populations.

In terms of genetic composition, multiple-merger coalescents translate into non-monotonic site frequency spectra (SFS). These non-monotonic SFS have already been observed in the context of Muller’s ratchet in [Neher and Hallatschek, 2012, Fig. 5B] and [Good et al., 2014]. Our findings go beyond this observations by providing direct access to the ancestry, allowing us to quantify the distortion of the genealogies as the strength of selection varies. Furthermore, our results suggest that the critical parameter for the onset of the Muller’s ratchet identified in [Etheridge et al., 2009] accurately predicts the emergence of multiple mergers in the associated genealogical structure.

While the potential similarity between the signatures of background selection and genetic hitchhiking was extensively investigated in [Schrider, 2020, Stephan, 2010], where it was shown to generate comparable genetic patterns, an analogous comparison between Muller’s ratchet and selective sweeps is still lacking. In this work, we conjecture that the similarity between the signatures of the latter two processes are even deeper, as they may produce identical genealogies. A statistical analysis of the power of the existing tests for selection to distinguish Muller’s ratchet and selective sweeps is needed to make further conclusions.

## 2 Model and Methods

### 2.1 The Wright–Fisher model with selection and mutation

In this work, we focus on a simple model for purifying selection, that is, the Wright–Fisher model with selection and mutation. We consider a finite haploid population of constant size *N*. All mutations have the same fitness cost *s* < 0, and the fitness of an individual with *k* mutations is given by (1 + *s*)^*k*^, i.e. selection is multiplicative. The parameter *s* will be referred to as the *selection coefficient*. We assume that the number of sites is effectively infinite, so that each mutation occurs at a new site and there are no back mutations.

The population evolves in discrete generations according to the standard Wright–Fisher dynamics. At generation *t* + 1, each individual picks a parent from generation *t* with probability proportional to the parental fitness. The individuals inherit the mutations of their parents. Each individual then acquires a Poisson number of new mutations with parameter *µ*. The parameter *µ* of this Poisson random variable represents the *genomic mutation rate*. In the remainder of this paper, we will write

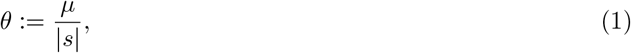

for the *mutation-selection ratio* and we will often refer to |*s*| as the *strength of selection*.

In Section 2.4, we develop statistics to characterise the genealogy of the Wright-Fisher model for a wide range of parameters (*µ, s*). These tools rely on coalescent theory, from which we recall some necessary facts in Section 2.2 and Section 2.3. The results of our simulations are presented in Section 3. Finally, we compare our conclusions with previous work on both purifying selection and adaption in the discussion (Section 4).

### 2.2 Coalescent theory

The universality class of Beta-coalescents [Pitman, 1999, Sagitov, 1999] is a one-parameter family of genealogies that can be seen as a generalisation of the neutral coalescent developed by [Kingman, 1982]. For *α* ∈ (0, 2), the coalescence rates of the Beta(2 − *α, α*)-coalescent are given by

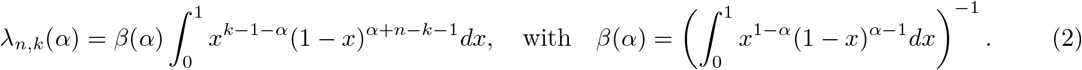

More precisely, *λ*_*n,k*_(*α*) corresponds to the rate at which the first *k* out of *n* ancestral lineages coalesce into a single lineage. Note that the Bolthausen-Sznitman coalescent is a Beta(1, 1)-coalescent (*α* = 1) and that Kingman’s coalescent can be recovered by letting *α* → 2. In the latter case, the rates (2) become

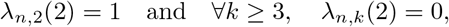

and the coalescent process only comprises binary mergers.

Heuristically, the parameter *α* controls the number and the size of the multiple mergers (*k* > 2) in the ancestry: we see from (2) that the probability to observe large coalescence events (*k* ≫1) increases as *α* decreases to zero. Hence, Beta-coalescents get further away from the neutral (binary) genealogy as *α* gets smaller (see Figure 1). For this reason, we will refer to *α* as the *distortion parameter*. We will say that small values of *α* correspond to large distortions of the genealogy (compared to neutrality, *α* = 2).

**Figure 1:**
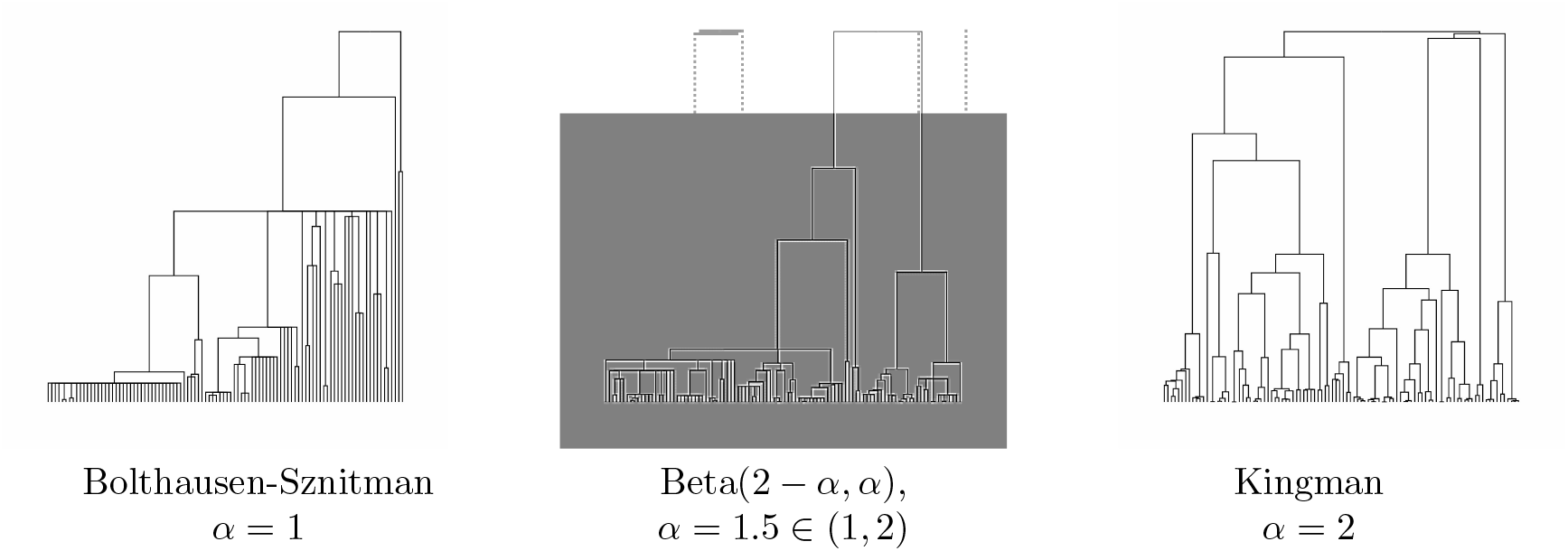
A simulation of Beta(2 *α, α*)-coalescents for *α* ∈ {1, 1.5, 2}. For *α* = 2 (Kingman), the genealogical tree is binary. In this figure, the three trees are scaled in such a way that they have the same height.

Beta-coalescents arise naturally in a wide range of population models, including models with selection [Berestycki et al., 2013, Schweinsberg, 2017, Cortines and Mallein, 2018] and spatial structure [Birzu et al., 2021]. Neutral genealogies (*α* = 2) are usually observed in populations with finite reproductive variance while distorted ones (*α* < 2) are obtained from highly skewed offspring distributions [Schweinsberg, 2003, Huillet and Möhle, 2021]. It was also shown in [Steinrücken et al., 2013] that the predictions derived from beta-genealogies fit the genetic data obtained from different species. Based on these observations, it is reasonable to assume that the genealogy in the Wright-Fisher model with mutation and selection also belongs to the universality class of Beta-coalescents.

### 2.3 Genetic diversity in beta-genealogies

The distortion parameter *α* has a significant impact on the patterns of sampled genetic variation. Not only does this parameter affect the shape of the ancestries but also the depth of the trees. Here, we give an overview of various statistics derived from beta-genealogies to show how these distortions in the genealogy translate in terms of genetic diversity.

#### Site frequency spectrum (SFS)

Let *p*_*α*_(*x*) be the site frequency spectrum, i.e., the number of mutations with frequency *x* ∈ (0, 1) in the population. It is known that for *α* = 2, the spectrum *p*_2_ is a monotonic function. More precisely, we have

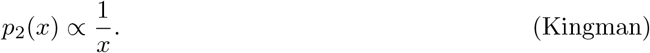

For *α* < 2, an explicit formula for *p*_*α*_ for intermediate frequencies is still lacking. For *α* = 1, asymptotics for small and large frequencies were derived in [Berestycki and Berestycki, 2014]:

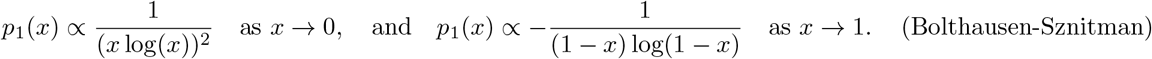

For *α* ∈ (1, 2), these read [Okada and Hallatschek, 2021]

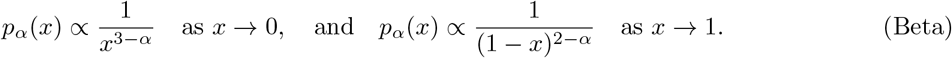

In particular, these formulas show that the SFS is no longer monotonic when *α* < 2. We also see that a decrease in *α* results in an increase in the expected number of very rare alleles.

#### Fixation probability

Let us now discuss the fate of a single mutant: assume that the population now consists of *N* − 1 individuals with fitness 1 (wild type) and a single mutant with fitness 1 + *s*. In this paragraph, we assume that *s* > 0 (positive selection) and that the strength of selection goes to zero as the population size *N* increases. Under suitable conditions, it was shown (see e.g. [Haldane, 1927, Birkner et al., 2023, Okada and Hallatschek, 2021]) that the probability *p*_*fix*_ for the mutant allele to become fixed is given by

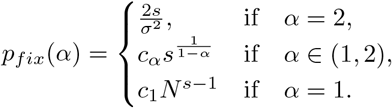

These formulas indicate drastic differences in the behaviour of the allelic trajectories.

#### Depth of the tree

Intuitively, coalescence events involving a large number of ancestral lineages generate shallow ancestries. Thus, we see from Section 2.2 that the natural timescale of the evolution decreases as *α* decreases. Typically, Kingman’s coalescent is observed on a timescale that corresponds to the total population size *N* or to the *effective* population size *N*_*e*_. In the latter case, this indicates that not all individuals take part in the evolutionary dynamics. For *α* ∈ (1, 2), the coalescent process usually evolves on a polynomial or polylogarithmic timescale [Huillet and Möhle, 2021].

### 2.4 The distortion estimator

In this work, we aim to estimate the distortion parameter *α* from simulations of the Wright-Fisher dynamics. We adapt the approach developed in [Brunet et al., 2006, Brunet and Derrida, 2012] based on coalescence times and extend it to beta-genealogies.

Let *T*_*k*_ denote the time to the most recent common ancestor of a sample of size *k* in the Wright–Fisher model and write 𝔼 [*T*_*k*_] for the corresponding expectation. Clearly, the expected depth of the tree scales like 𝔼 [*T*_2_] (or equivalently, like 𝔼 [*T*_*k*_], *k* > 2). On the other hand, the shape of the tree can be characterised through the *coalescence ratios*

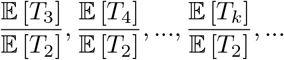

Indeed, we see that coalescence events involving a large number of ancestral lineages decrease the coalescence ratios. Thus, it is reasonable to believe that there exists a one-to-one relation between these ratios and the distortion parameter *α*.

In fact, one can check (we refer to Appendix A.2 for a proof of this result) that the coalescence ratios of the Beta(2 − *α, α*)-coalescents satisfy

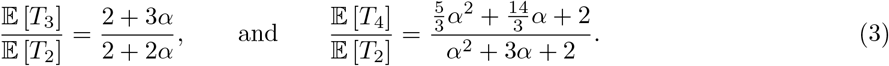

In particular, we recover the values from [Brunet et al., 2006] for *α* = 0, 1, 2. Note that one could go on calculating the ratios for *k* ≥ 5 (the calculations are similar to that derived in Appendix A.2). Here, we will only use the two identities given in (3) and verify that they provide the same value for *α*. The main advantage of this method lies in the fact that the coalescence ratios are scale-free and thus allow us to work directly on the tree structures without any *a priori* knowledge on the natural timescale of the evolution.

In practice, we will estimate the distortion parameter using the average coalescence time ⟨*T*_2_⟩, ⟨*T*_3_⟩ and ⟨*T*_4_⟩ obtained from simulations of the Wright–Fisher model. Inverting (3), we obtain the following two unbiased estimators

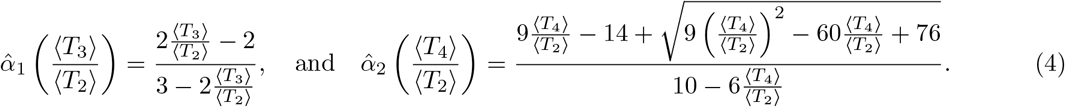

We conclude this section by remarking that the coalescence times *T*_1_, *T*_2_, … computed from different samples of the same genealogical tree are correlated. In our simulations, we will average these coalescence times on several realisations of the Wright–Fisher dynamics, i.e. on several realisations of the genealogy. This will ensure that our estimators 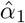 and 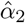 are consistent.

### 2.5 Simulations

In our simulations, we compute the average coalescence times ⟨*T*_2_⟩, ⟨*T*_3_⟩ and ⟨*T*_4_⟩ from independent iterations of the Wright-Fisher dynamics. In order to keep track of these quantities, we record the index of the parent of each individual in every generation. For each iteration of the Wright-Fisher model, we reconstruct the full genealogy by tracing back the ancestral lineages from the final generation *t*_*f*_. The quantity ⟨*T*_*k*_⟩ is then computed as follows: we sample *k* individuals uniformly at random from the genealogy and calculate the time to their most recent common ancestor (MRCA). We repeat this procedure *n* times in one genealogy and average this quantity across *m* independent iterations of the Wright–Fisher dynamics.

In Section 3, we present the results of these simulations for *n* = *N* = 3000, *m* = 200 and running time *t*_*f*_ = 5*N* generations. The running time *t*_*f*_ is chosen in such a way that, with high probability, the *N* individuals alive at *t*_*f*_ have found their MRCA. We expect this time to be of order 2*N* for Kingman’s coalescent and to be much smaller than *N* in beta-genealogies, see Section 2.3. Moreover, this choice of *t*_*f*_ ensures that the average coalescence times ⟨*T*_*k*_⟩ are independent of the initial condition for the fitness distribution. The values for *µ* range from 0.01 to 0.3 and for *s* from 0 to −0.15. For *s* = 0, the Wright–Fisher model is neutral.

The simulations were performed in Python on the *ISTA High-Performance Computing Cluster* and the code is available at https://github.com/khudyakovaks/genealogy_fitness_waves.

## 3 Results

### 3.1 The phase diagram

This section is dedicated to the results of the simulations presented in Section 2.4 and Section 2.5. As we shall see, these simulations uncover a rich phase diagram.

#### Distortion estimators

First, we observe that the estimators 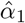 and 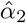 provide similar values for the distortion parameter *α*, see Figure 2. This supports the assumption that the Wright-Fisher model with selection and mutation belongs to the universality class of Beta-coalescents. For this reason, we will restrict our analysis to statistics obtained from ⟨*T*_2_⟩ and ⟨*T*_3_⟩. We refer to Appendix A.1 for additional material on the statistics obtained from ⟨*T*_4_⟩.

**Figure 2:**
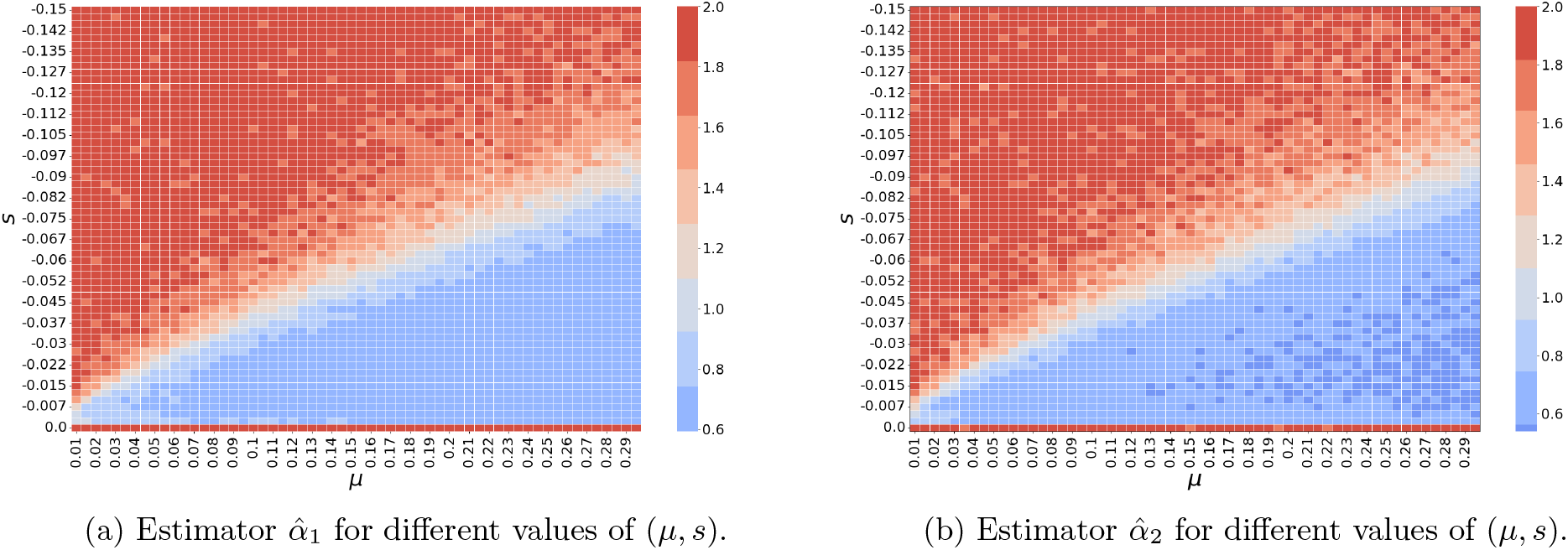
Estimators 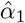 and 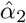 for different values of (*µ, s*). Each square represents the average value of the distortion estimators obtained from *m* = 200 simulations of the Wright-Fisher dynamics after *t*_*f*_ = 5*N* generations. Recall from Section 2.4 that we set *N* = 3000.

As the strength of selection and mutation rate vary, we observe a wide range of values for the distortion estimator 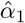 (see Figure 2). When the strength of selection |*s*| is large compared to the mutation rate *µ* (as in the top left corner of Figure 2), the estimator 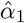 is close to the distortion parameter corresponding to Kingman’s coalescent 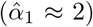 Similarly, on the horizontal line *s* = 0, the distortion estimator is close to *α* = 2, which corresponds to the parameter associated to Kingman’s coalescent. For fixed *µ* (vertical lines), as the selection parameter increases to 0, the distortion estimator 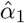) first remains broadly constant equal to 2, ii) then decreases to some level *α*^*^(*µ*) *<* 2, iii) remains at this level iv) and finally abruptly increases to *α* = 2 as the selection parameter *s* tends to 0. This non-monotonic behaviour is illustrated in Figure 3 (b). Recall from Section 2.2 that *α* < 2 indicates the presence of multiple mergers in the underlying genealogical structure. The above discussion then suggests the following phase diagram:

**Figure 3:**
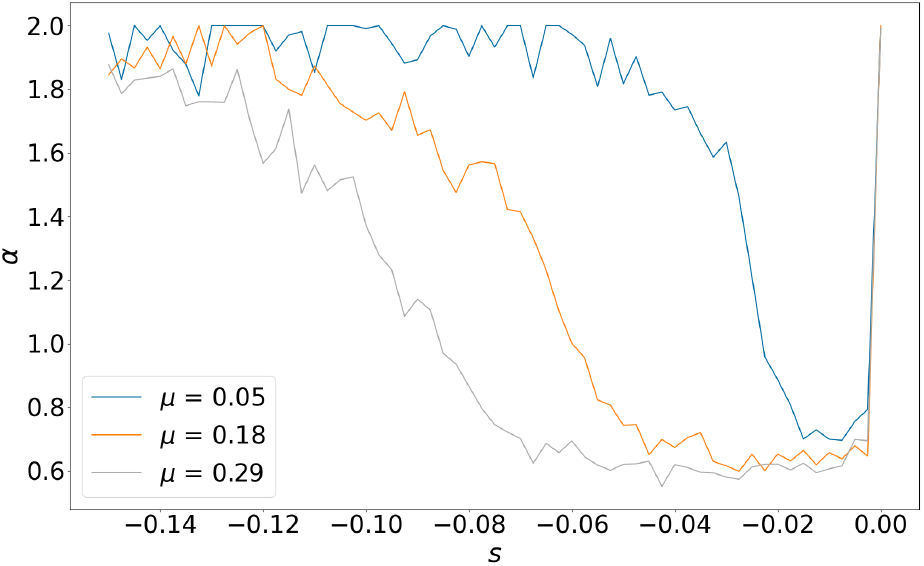
Estimator 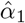 as a function of *s* for *µ* ∈ {0.05, 0.18, 0.29}. Each line represents the estimator 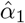 as a function of *s* for a fixed value of *µ* (vertical lines on Figure 2 (a)).

- The set of parameters (*µ, s*) such that *α*(*µ, s*) = 2 corresponds to the **Kingman regime**. This regime coincides with the top left red triangle in Figure 2,
- The set of parameters (*µ, s*) such that *α*(*µ, s*) *<* 2 corresponds to the **multiple-merger regime**. In Figure 2, this regime covers the blue and white regions (bottom right triangle).

#### Depth of the ancestry

We recall from Section 2.4 that the estimators 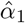 and 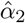 are independent of the height of the genealogical trees and thus only allow us to characterise the shape of the trees. To complement this picture, we need to define a statistic characterising the depth of the beta-genealogies identified in Figure 2. We also recall from Section 2.4 that the mean height of these genealogies is proportional to the mean pairwise coalescence time 𝔼 [*T*_2_].

For the classical Wright-Fisher model, i.e. for *s* = 0, we know that 𝔼 [*T*_2_] = *N* [Kingman, 1982]. In sharp contrast, multiple-merger coalescents have proved to emerge on much shorter timescales, see Section 2.3. For instance, in the models of waves of species range expansion [Birzu et al., 2021, Tourniaire, 2021], this scaling *s*_*N,α*_ only depends on *α* and *N* and is given by

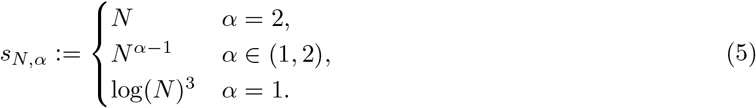

In view of the above discussion, we expect to observe a decrease in the height of the genealogies as the parameters (*µ, s*) enter the multiple-merger regime. To test this, we plot the estimator

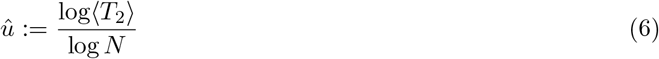

in Figure 4 (b). As we shall see, this estimator is well-adapted to capture polynomial scalings.

**Figure 4:**
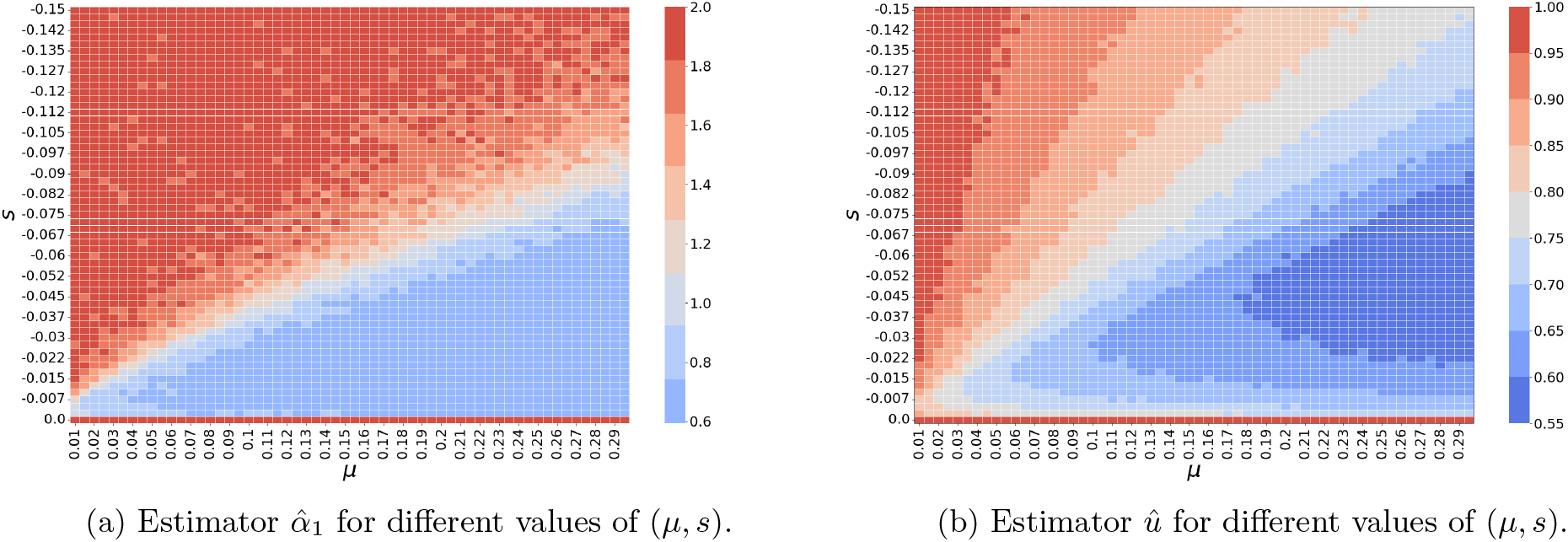
Estimators 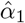 and 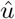 for different values of (*µ, s*). We see that, in Kingman regime, 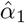 is roughly constant equal to 2 (upper triangle in the LHS figure) but the depth of the genealogy varies (same region in the RHS figure).

Interestingly, unlike spatial waves (eq. (5)), we observe a decrease in the depth of the ancestry within the Kingman regime (*α* = 2). Our simulations suggest that when 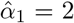, the genealogy is given by Kingman’s coalescent on a timescale of order 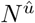, where the exponent 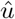 is non-decreasing with respect to the slopes of the lines |*s* | */µ*, see Figure 4 (b). In other words, in the Kingman regime, the evolution is indistinguishable from a neutral evolution with a reduced effective population size 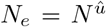. This is in line with previous work [Walczak et al., 2012, Nicolaisen and Desai, 2012] and will be discussed later in this section.

### 3.2 Muller’s ratchet

Previous work on wave-like phenomena in evolving populations has shown that the genealogical structure of a wave is closely related to its speed (see e.g [Brunet and Derrida, 2012] in the context of adaptation and [Birzu et al., 2021] for spatial waves). In this section, we investigate the existence of a similar connection for purifying selection.

#### The rate of the ratchet

The evolution of the fitness profile of an asexual population can be modelled as the propagation of a *fitness wave* travelling in the space of fitness with constant speed and shape. Depending on the values of the parameters (*µ, s*), if *s* < 0, the wavefront is either static or advancing to lower fitness levels. The latter phenomenon is known as *Muller’s ratchet* and the speed of the wave is referred to as the *rate of the ratchet* [Haigh, 1978]. In the absence of back mutations, the best-fit class (i.e. the least-loaded class) can go extinct: we say that the ratchet *clicks*.

The following rule of thumb was derived in [Etheridge et al., 2009] to predict the rate of the ratchet. The authors identified the quantity

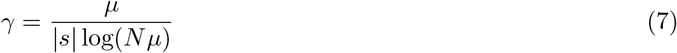

as the key parameter: the expected time between two clicks is of the order

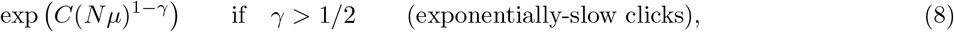

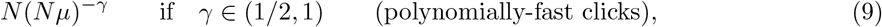

as long as *Nµ* ≤ 900; see [Etheridge et al., 2009, Section 5]. Note that, in our simulations, we have 30 ≤ *Nµ* ≤ 900 so that this rule can be applied without modifications. When *γ* > 1 and *Ne*^−*θ*^ *<* 1 (see (1)), the clicks are much more frequent than in the polynomially-fast regime.

In practice, the exponentially-slow clicks are too slow to be observed in real populations. The population thus achieves a *stable* (or *deterministic*) mutation-selection balance. In sharp contrast, for *γ* > 1/2, on the timescale of the evolution, the system fails to achieve mutation-selection balance and we say that the ratchet operates. This phenomenon is illustrated in Figure 5.

**Figure 5:**
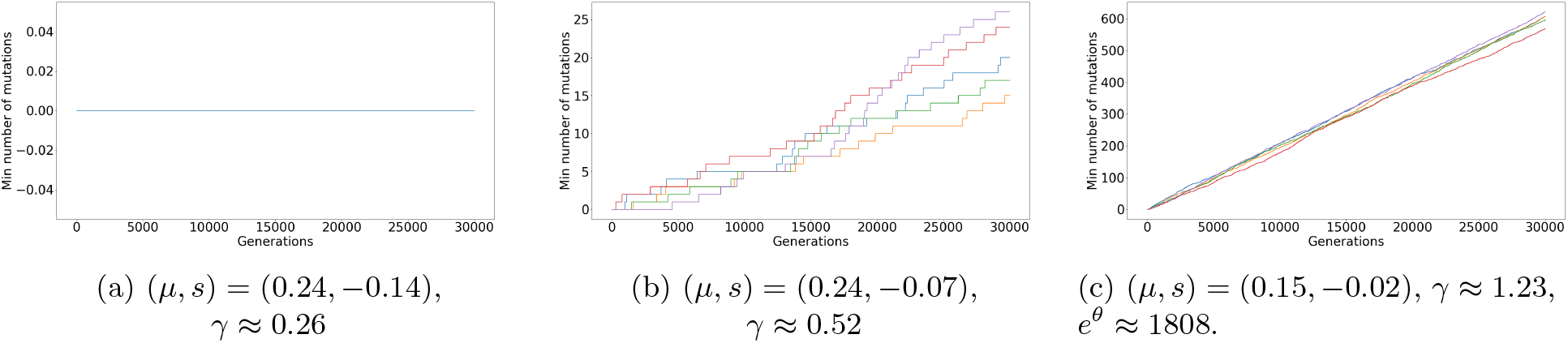
Load in the best class for five simulations of the Wright-Fisher dynamic spanning the different regimes of the ratchet. For each pair of parameter, we plot the number of mutations present in the fittest class for *N* = 3000 and 10*N* generations.

#### The critical line

We now compare these different regimes for the rate of the ratchet with the results of our numerical simulations, see Figure 6.

**Figure 6:**
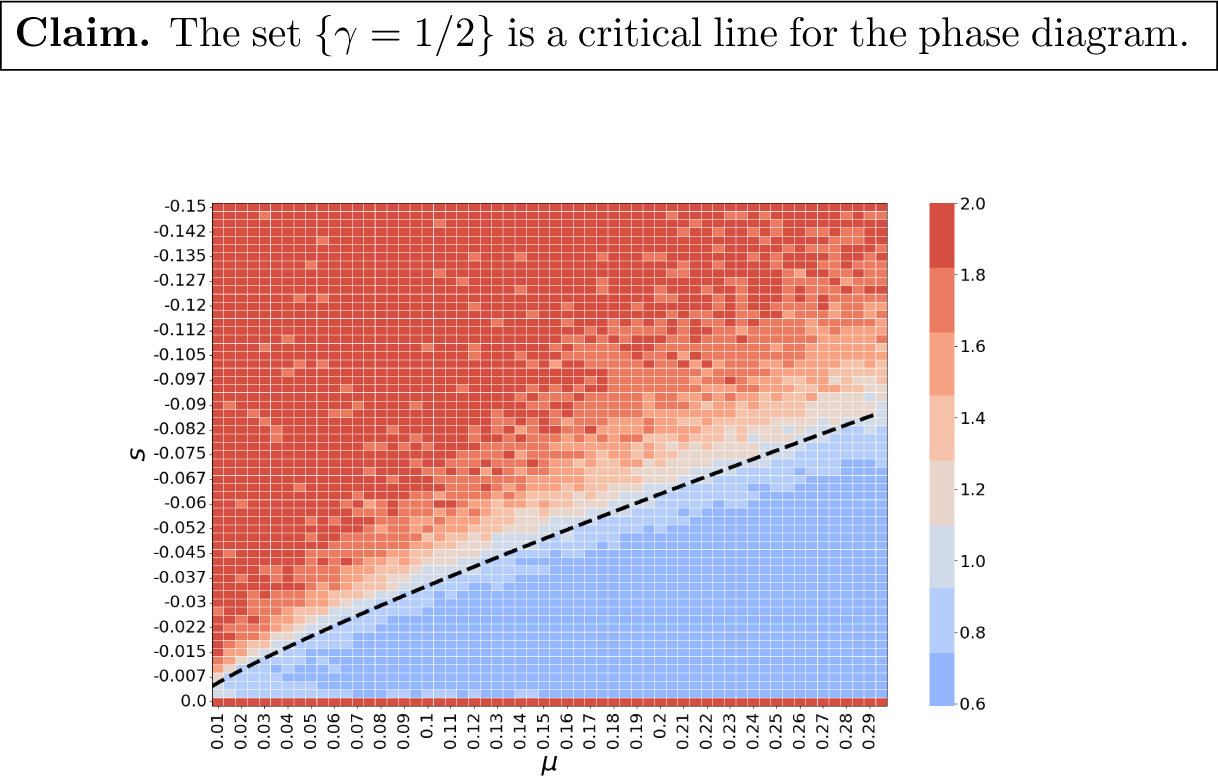
Estimator 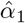 for different values of (*µ, s*) and the critical line {*γ* = 1/2}.

It becomes clear from Figure 6 that the set {*γ* = 1/2} is a critical line for the phase diagram introduced in Section 3.1. Indeed, we observe that the transition between the Kingman regime 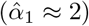 and the multiple-merger regime 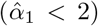 occurs precisely on the {*γ* = ½} line. In words, this suggests that Kingman’s coalescent is the universal genealogy of populations achieving a stable mutation-selection balance. As soon as the ratchet starts operating, the genealogical tree deviates from the neutral genealogy: the ancestry is described by a multiple-merger coalescent.

### 3.3 Heuristics

In this section, we develop heuristics to support our claim.

We consider the following parametrisation of the mutation rate and of the selection parameter:

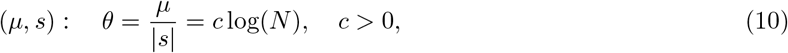

which amounts to considering lines with slopes (*c* log(*N*))^−1^ on the two-dimensional plots. This scaling appears naturally in the *near-critical* regime [Igelbrink et al., 2024] in which *µ* scales like a power of *N* (*µ* = *N* ^−*m*^ with *m* ∈ (0, 1)) and *s* is chosen in such a way that *γ* stays constant as *N* increases. By definition of *γ* (see (7)), it is sufficient to set

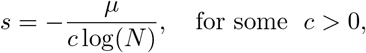

which coincides with (10).

#### The Kingman regime

If the population achieves a stable mutation-selection balance, the fitness wave is well approximated by a Poisson distribution whose parameter is given by the mutation-selection ratio *θ* [Haigh, 1978]. Hence, under (10), the expected number of individuals with *k* mutations is approximately given by

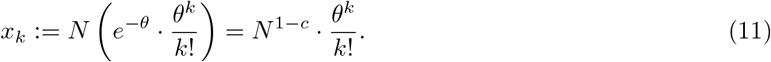

Sample *n* individuals from the population and denote by *k*_1_, …, *k*_*n*_ their loads. A Chernoff bound shows that, for all *i* ∈ {1, …, *n*} and *k > θ*,

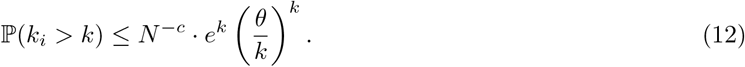

In particular, we see that, with high probability

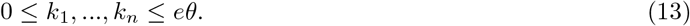

It then follows from Stirling’s formula 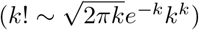 that

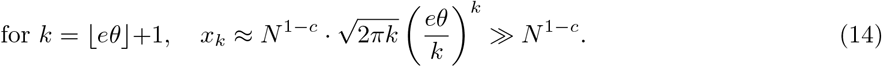

Putting all of this together, we obtain that, with high probability,

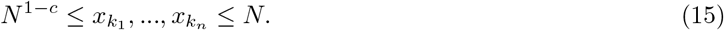

We now trace back the *n* lineages corresponding to these *n* individuals. Backward-in-time, the lineages evolve as follows:

i. *mutation events:* each lineage in class *k* looses one mutation at rate *sk*, see [Walczak et al., 2012].
ii. *coalescent events:* two lineages can coalesce if they belong to the same class. The probability to observe a coalescence event is inversely proportional to the size of the class (“neutral evolution within the classes”).

Hence, along one lineage, the expected time between two mutation events is bounded above by 1/ |*s* |. Moreover, note that (15) holds at each time step since the load of a lineage can only decrease. Thus, the expected time between two coalescence events is bounded below by *N* ^1−*c*^. On the other hand, the condition *γ* < ½ can be rewritten as

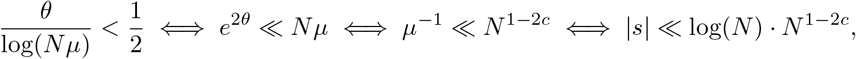

see (10). Multiplying the upper bound for the expected time between two mutation events by the maximal distance an individual has to travel to reach the zero-loaded class (see (13)), we get that

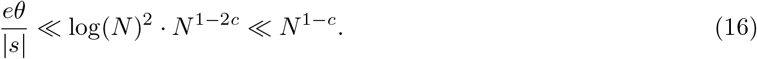

Therefore, in the large-*N* limit, we do not observe any coalescence event before all the lineages reach the first class. The backward-in-time dynamics is thus divided into two consecutive stages: the *travelling phase* (the lineages travel back to the zero-loaded class) and the *coalescing phase* (they merge until a single one remains). We see from (16) that the duration of the travelling phase is negligible compared to that of the coalescing phase. As a consequence, the limiting genealogy only depends on the dynamics of the lineages in the coalescing phase. This phenomenon, akin to the “collapse of structure” studied in [Nordborg and Krone, 2002] for spatially structured populations, is illustrated in Figure 7.

**Figure 7:**
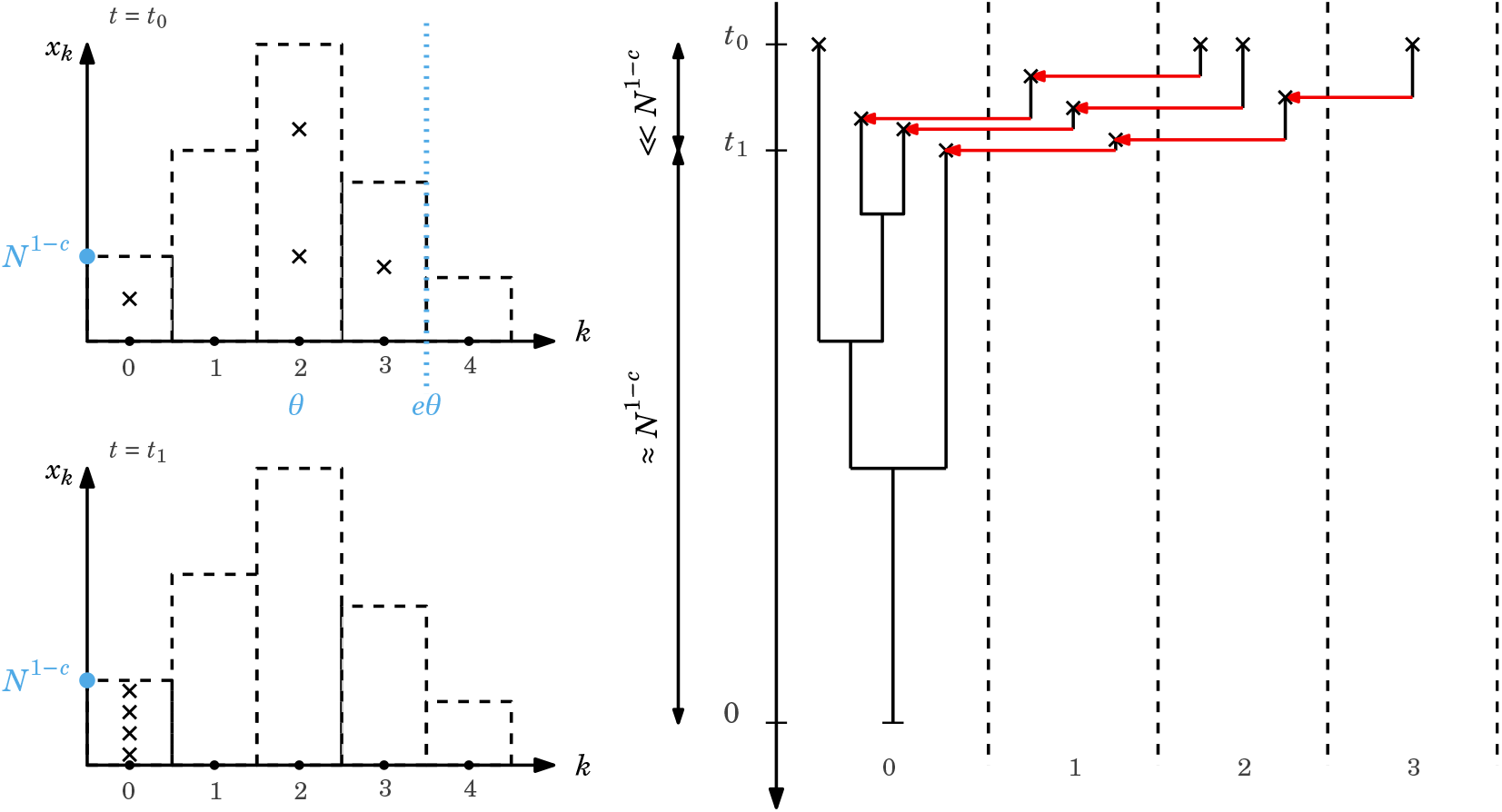
Backward-in-time dynamics in the Kingman regime. We sample *n* = 4 individuals at time *t*_0_. On the top left figure, the sampled individuals are represented by crosses. With high probability, these individuals are sampled to the left of the line *k* = *eθ* in the Poisson distribution (blue vertical dotted line). Their loads are respectively *k*_1_ = 0, *k*_2_ = *k*_4_ = 2 and *k*_3_ = 3. The dynamics of the corresponding lineages are represented on the right figure. The lineages first travel back to the zero-loaded class (travelling phase). At time *t*_1_, the *n* lineages are in the zero-loaded class. With high probability, there is no coalescence event between times *t*_0_ and *t*_1_. The lineages then start coalescing. Since they all carry the same number of mutations (*k* = 0), they behave as in a neutral population with size *x*_0_ = *N* ^1−*c*^. The genealogy of the sample is thus given by Kingman’s coalescent on the timescale *N* ^1−*c*^ started at time *t*_1_. The duration of the travelling phase *t*_0_ − *t*_1_ will be referred to as the *delay D*.

These heuristics are consistent with the results of our simulations. For large *N*, the travelling phase is nearly imperceptible. Moreover, in the coalescing phase, the dynamics of the lineages is that of a neutral genealogy. As a result, the limiting genealogy is given by Kingman’s coalescent, as observed in Figure 6.

It follows from the above discussion that the timescale of the genealogy should correspond to the expected size of the zero-loaded class (*N* ^1−*c*^, see (11)). In our setting, we see that the exponent in this polynomial timescale is given by the slope of the (*µ, s*)-lines introduced in (10). This connection also appears in Figure 8: above the critical line, the estimator for the exponent 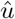 only depends on the mutation-selection ratio *θ*. Finally, we remark that, for finite *N*, the travelling phase translates into a *delay* at the leaves of the genealogical tree associated to the sample (see Figure 7). This delay *D* should be well approximated by the expected time it takes for a lineage to travel back from the mode of the Poisson distribution (≈ *θ*),

**Figure 8:**
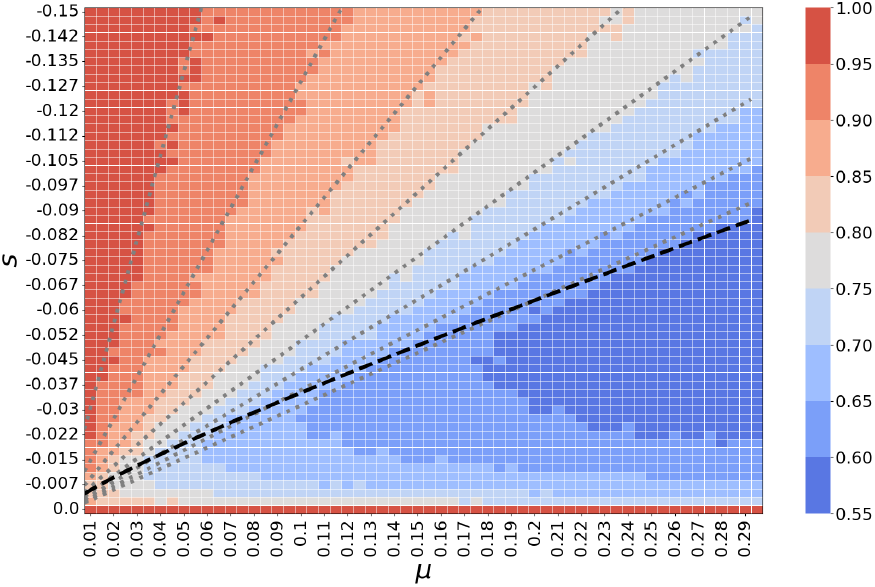
Estimators 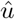 for different values of (*µ, s*); see (6). The dashed line corresponds to the critical line {*γ* = 1/2} and the dotted lines to the *c*-lines parametrised by (10).

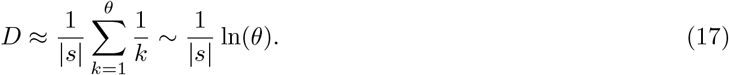

We see that the delay *D* increases as the strength of selection decreases. This observation explains the noise above the critical line in Figure 6. Indeed, we see from the above heuristics that the shape of the beta-genealogy only depends on the dynamics in the least-loaded class. Therefore, the distortion parameter *α* should be derived from the coalescence times within the best class. Yet, the coalescence time ⟨*T*_*k*_⟩ is the sum of the delay *D* and the average coalescence time within the best class. Moreover, one can check that 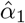 and 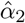 are decreasing functions of the coalescence ratios and that *a/b* > (*a* + *D*)/(*b* + *D*) for all *a, b, D* > 0. This shows that we tend to underestimate the distortion *α* when the delay *D* is comparable to *N* ^1−*c*^. In the simulations, this is the case when the population size *N* is small or when the strength of selection is small compared to the mutation rate.

#### The multiple-merger regime

From Section 3.2 and (10) we know that for 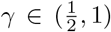, the expected interclick times are of the order *N* ^1−*c*^. In other words, the fluctuations in the class sizes develop on the same time scale as the neutral genealogies in the best class (see Figure 7). The above heuristics thus break down. In the Wright-Fisher (forward-in-time) dynamics, before a click happens, the least-loaded class is much smaller than its expected size *x*_0_. In the backward-in-time picture, this size reduction may give rise to an accumulation of binary mergers. Indeed, if more than two lineages jump back to the new best class before it reaches a macroscopic size (i.e. of order *N* ^1−*c*^), they will merge much faster than in the Poisson approximation, see Figure 7 and Figure 9. On the evolutionary timescale, this accumulation of binary mergers in the reduced best class is indistinguishable from a multiple merger, see Figure 9.

**Figure 9:**
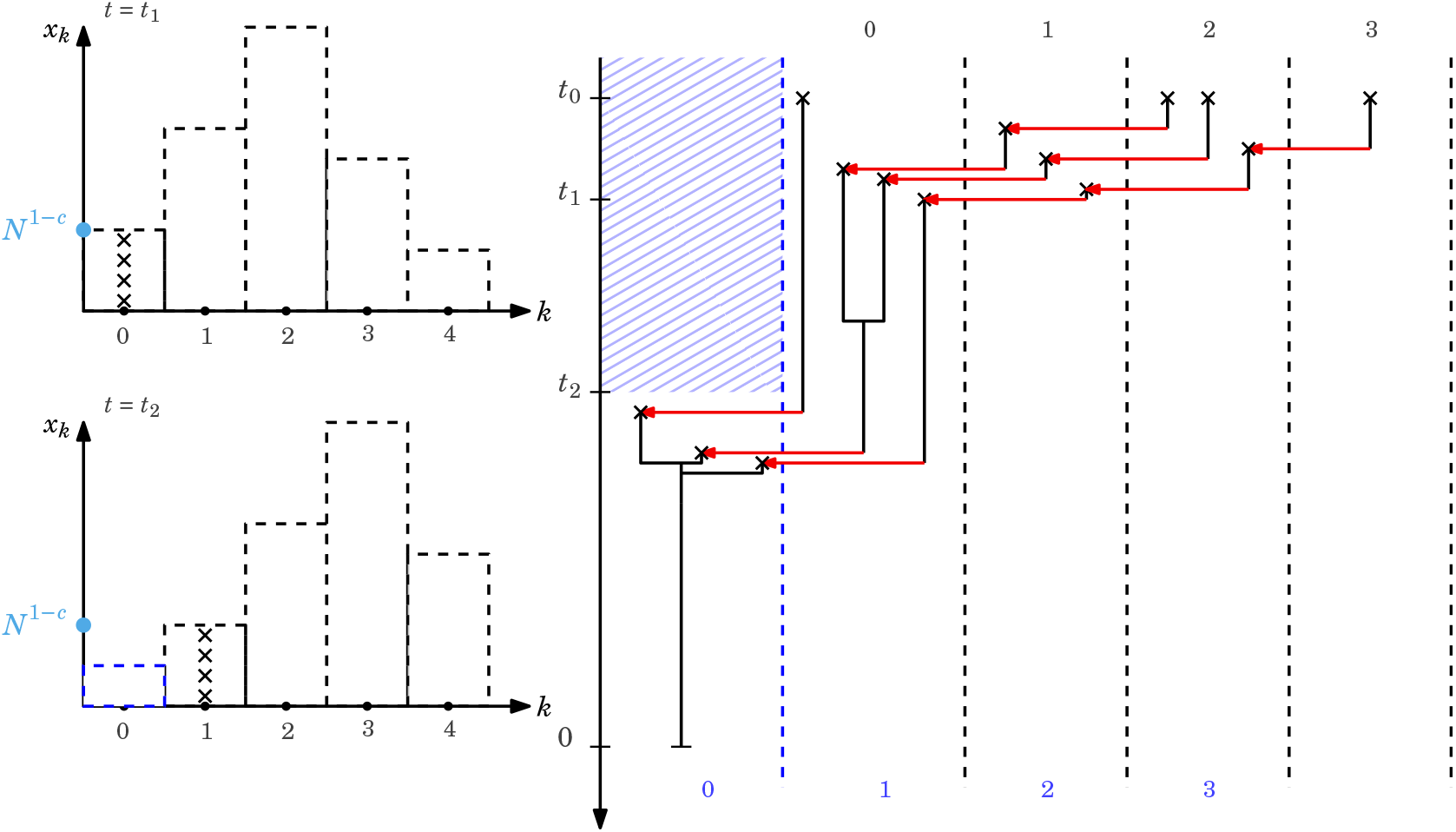
The multiple-merger regime for *γ* ∈ (1/2, 1). As in Figure 7, the lineages first travel back to the current least-loaded class. They then start coalescing in this class. At time *t*_2_, the ratchet clicks. All the lineages travel back to the new best class in a time of order *µ*^−1^. Multiple mergers appear in the limiting genealogy when the size of the best class is much smaller than *x*_0_.

When *γ* > 1, we see from (15) that *x*_0_ ≪ 1. Hence, the Poisson approximation indicates that, most of the time, the best class is empty. As observed by [Gessler, 1995], this causes a reduction of the variance in the fitness distribution (compared to the Poisson distribution) and large discrepancies in fitness values. We believe that these discrepancies translate into an explosion of the variance in reproductive success. Yet, we know from [Schweinsberg, 2003] that multiple-merger coalescents emerge in populations with infinite reproductive variance. Let us mention that analogous arguments were used in [Berestycki et al., 2013, Tourniaire, 2021, Foutel-Rodier et al., 2024] to characterise the genealogy of spatial waves.

The characteristics of the different regimes are summarised in Figure 10.

**Figure 10:**
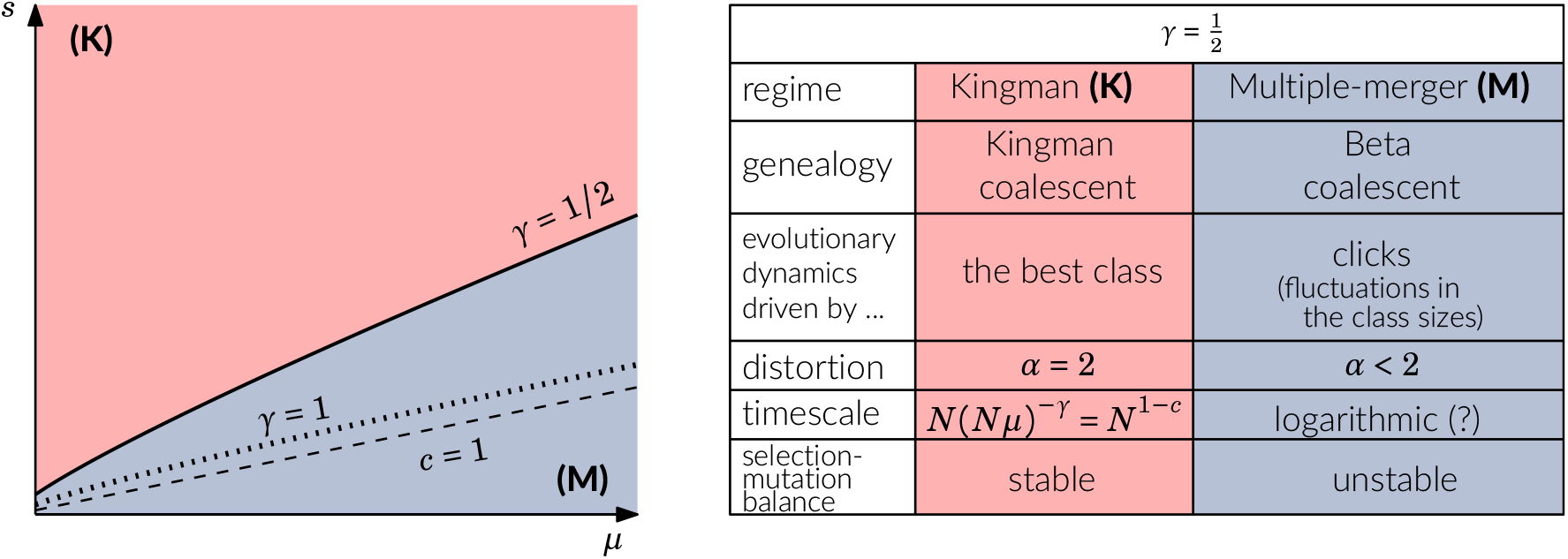
Characteristics of the two regimes.

### 3.4 Connection with previous theoretical work

The mathematical analysis of the Wright-Fisher dynamics turns out to be cumbersome. The constant population size and the selection mechanism induces strong dependencies between individuals. As a result, one can hardly identify the main evolutionary forces with no *a priori* knowledge on the (*µ, s*)-scaling.

In [Boenkost et al., 2024], we define a tractable model for selection against deleterious mutations. In this framework, we consider a critical branching particle system. The particles carry loads *y* evolving according to independent diffusions with positive drifts. The multiplicative selection is replaced by truncation selection: a particle is killed when its load becomes too high. This model relies on two postulates. First, we assume that the dynamics is driven by the best class: in this class, the effect of selection is negligible and the population behaves like a branching process. Hence, the limiting dynamics should not depend on the precise form of the selection mechanism. Note that this assumption is in line with the above heuristics. Second, the diffusion approximation hinges upon the fact that mutations have a small impact on fitness.

In the branching particle system, we are able to prove that the system settles in a stable configuration 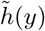 *dy* and derive the reproductive value *h*(*y*) of an individual with load *y* (the functions *h* and 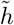 are explicit). In addition, we show that when the bulk of the stable load distribution 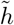 is located at a distance *c* log(*N*) of the origin (see (11)), the variance in the reproductive value

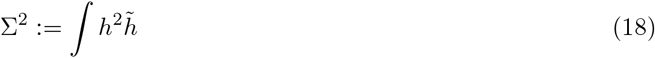

is of order *N*^*c*^ and the genealogy is given by Kingman’s coalescent on the timescale *N* ^1−*c*^. More precisely, we observe that all the coalescence events occur in a neighbourhood of the origin (0, *A*). One can then check, using the explicit formulae for *h* and 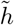, that the expected number of individuals whose load is smaller than *A* is of order *N* ^1−*c*^ and that the mass of the integral (18) is concentrated in (0, *A*). For these reasons, the interval (0, *A*) is interpreted as the *best class* in the continuous model.

Note that all the scalings obtained from the critical branching particle system are reminiscent of those observed in the Wright-Fisher dynamics in the stable mutation-selection balance (*γ* < 1/2). Moreover, our findings support the assumption that the genealogy of a structured population strongly depends on its reproductive variance.

## 4 Discussion

In this work, we observe a phase transition in the genealogy of the Wright–Fisher model with selection and mutation as the strength of selection varies. Our method allows us to directly observe the genealogy of a sample, without any *a priori* knowledge of the size of the fitness classes or the strength of selection. Additionally, we derive heuristics relating the transition in the dynamics of the ancestral lines to the onset of Muller’s ratchet in the Wright–Fisher model.

When selection is strong enough to maintain the zero-loaded class at the stable mutation-selection balance, the genealogy is described by a time-changed Kingman coalescent. This time change is determined by the effective population size of the population, which is reduced to the size of the fittest class. These observations are consistent with the theory of background selection [Charlesworth et al., 1993]. As the strength of selection decreases, the genealogies transition from Kingman’s coalescent to multiple-merger coalescents, with the distortion parameter decreasing as the rate of the ratchet increases. Furthermore, we observe that the shift from binary to multiple-merger genealogical structures coincides with the onset of the ratchet. Hence, we deduce that the critical parameter *γ* identified in [Etheridge et al., 2009] for the rate of the ratchet is also a critical parameter for the associated backward-in-time dynamics.

### Connection with previous work

Similar transitions were observed in a wide range of models for adaptive evolution, see for example [Brunet and Derrida, 2012, Cortines and Mallein, 2018]. Therein, a higher strength of selection is associated with a stronger deviation from Kingman’s coalescent, resulting in genealogies involving large multiple mergers. The behaviour that we observe in the case of deleterious mutations is richer: as the strength of purifying selection increases, at first, multiple mergers emerge, but as the selection strength reaches a point where it can hold the ratchet, the genealogies are back to Kingman’s coalescent with a reduced population size. Therefore, moderate purifying selection results in the most extreme genealogies, whereas the absence of selection or strong purifying selection results in neutral genealogies.

In the purifying selection case, there are several works deriving approximations for the statistics of the genetic diversity [Desai et al., 2012, Walczak et al., 2012, Strütt et al., 2024, Cvijović et al., 2018, Good et al., 2014]. Using these approximations, it is possible to make conclusions about the underlying genealogical structure that causes the observed distortions in the genetic diversity. Here, we take a complementary approach: we are able to predict the genealogy of a sample directly, which in turn can predict genetic diversity using the results of coalescent theory.

In [Cvijović et al., 2018], the authors study the properties of the site frequency spectrum of a neutral mutation linked to a site under purifying selection. They consider the regime of strong selection, with *N* |*s*| *e*^−*µ/*|*s*|^ ≫ 1. Using a forward-in-time modelling approach, they derive an approximation for the site frequency spectrum (SFS), and show that the SFS differs from the neutral prediction and is non-monotonic. A non-monotonic SFS for Beta-coalescents can only be observed for *α* < 2 (see Section 2.3), that is for multiple-merger beta-genealogies. At a first sight, their result contradicts ours: under strong selection, we only see Kingman genealogies. We believe the reason for this apparent contradiction is the following. We have found that the emergence of multiple mergers coincides with the onset of the Muller’s ratchet, predicted by the critical parameter *γ* = 1/2 [Etheridge et al., 2009]. According to our results, the condition for the strong selection regime *N* |*s* |*e*^−*µ/*|*s*|^ ≫ 1 considered in [Cvijović et al., 2018] seems less relevant: depending on the values of *µ* and *s* chosen, we can observe genealogies described either by Kingman’s coalescent or the Bolthausen-Sznitman coalescent, both while *N* |*s*| *e*^−*µ/*|*s*|^ ≫ 1. This might explain why [Cvijović et al., 2018] observe non-monotonic and monotonic SFSs, depending on the choice of the ratio *µ*/|*s*|. Furthermore, part of the distortion in the SFS observed in [Cvijović et al., 2018] can be explained by the delay at the leaves (see (17)). Indeed, the delay until ancestral lineages can coalesce results in an excess of rare mutations in the SFS compared to the neutral expectation, even when the limiting genealogy is neutral.

### Possible sources of biases

One has to keep in mind that delays might distort the estimated value of *α*, giving the possibility to wrongly detect multiple-merger genealogies. This phenomenon could occur when the genealogies get shallower, in particular this might happen at the critical line. Increasing the number of individuals in the simulations would improve the estimation of *α* and help to decrease the chance of misclassification.

### Implications for the tests for selection

The importance of linked selection on the patterns of genetic diversity has long been recognised [Maynard Smith and Haigh, 1974, Charlesworth et al., 1993]. Our results indicate that the reduction in the diversity due to linkage with sites under selection not only qualitatively similar between positively and negatively selected mutations, but should actually be completely identical in the regions where Muller’s ratchet operates, due to emergence of multiple-mergers coalescents. Furthermore, our findings suggest that the tests for selection that rely on the reduction in genetic diversity are not able to distinguish between selective sweeps and the operation of Muller’s ratchet. To make further conclusions about the expected levels of genetic diversity under purifying selection, one needs to study models incorporating recombination on linear genomes.

### Perspectives

Genealogies under purifying selection exhibit a rich phase diagram which is only partially understood. In particular, studying the multiple-merger regime raises challenging questions. This lays the groundwork for future projects aimed at fully characterising the beta-genealogies in this regime, thereby determining whether the transition between Kingman and Bolthausen-Sznitman coalescents consists in a continuum of Beta-coalescents or in a sharp discontinuity. Moreover, in Figure 2, we estimate values of *α* ∈ (0.6, 1) in the blue region. For Wright-Fisher models, beta-genealogies with distortion parameters smaller that 1 are usually not observed. Instead, the limiting genealogies are often given by discrete-time coalescents [Huillet and Möhle, 2021, Schweinsberg, 2003], which do not belong to the class of Beta-coalescents. Here, it might be reasonable to believe that the values of *α* < 1 are due to an underestimation of the distortion parameter. As |*s*| decreases, the delay (17) increases and becomes particularly pronounced as the mean coalescence time decreases. This phenomenon comes into play in the multiple-merger regime, making it complex to come to a comprehensive understanding of the phase diagram.

## Data availability

The code used to perform the individual-based simulations is available at https://github.com/khudyakovaks/genealogy_fitness_waves.

## Author contributions

All three authors contributed equally to this work.

## A Appendix

### A.1 Additional material for Section 3

#### Rate of the ratchet

Our claim from Section 3.2 states that the shift from binary (*α* = 2) to multiple-merger (*α* < 1) genealogies coincides with the onset of the ratchet. We also recall from Section 3.2 that the onset of the ratchet is well predicted by the critical line {*γ* = 1/2}; see (7). In [Gessler, 1995], it is observed that a second transition in ratchet rate occurs at {*γ* = 1} : for *γ* > 1, clicks are more frequent than the polynomial rates given by (9). A natural follow-up question is whether this second critical line also plays a role in the phase diagram of the genealogies. In Figure 11, we plot these two lines on top of the heat maps obtained for our estimators 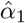 and 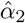. We observe that, while the {*γ* = ½} -line accurately predicts the emergence of multiple mergers, the {*γ* = 1} -line does not seem to correspond to any noticeable change in the structure of the genealogies.

**Figure 11:**
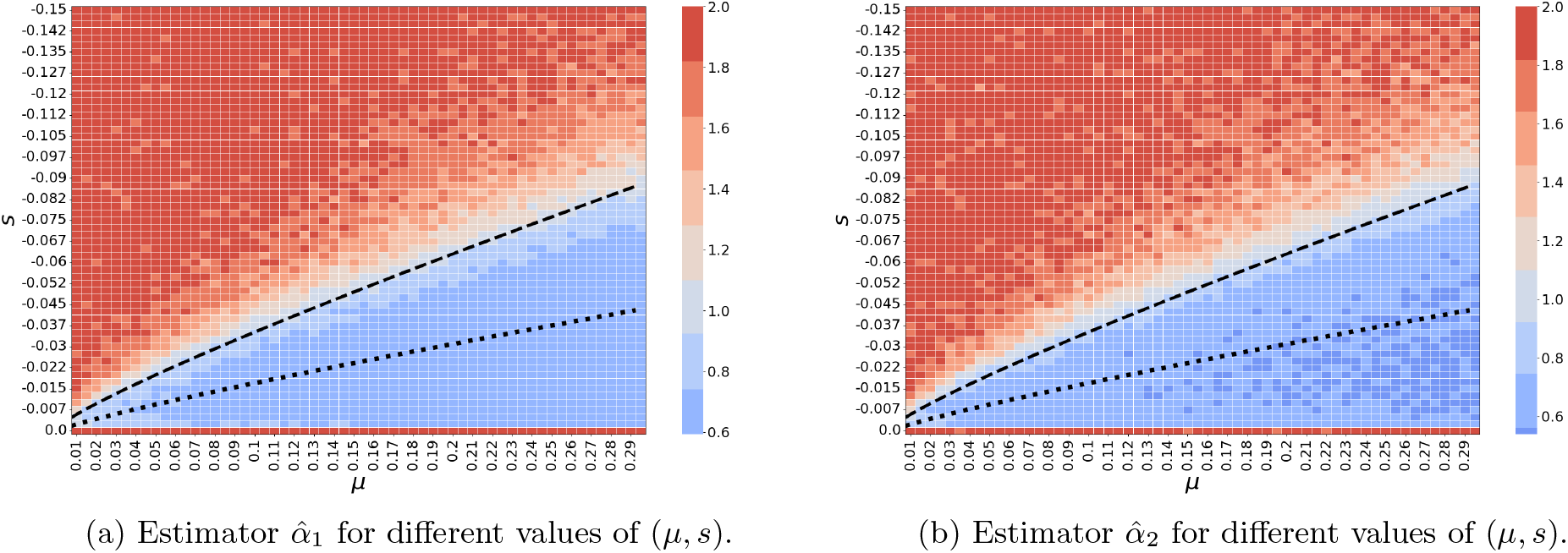
Distortion estimators for different values of (*µ, s*). On both figures, the dashed line correspond to {*γ* = 1/2} and the dotted line to {*γ* = 1}.

We repeat the same procedure for the estimator 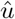 (see (6)) on Figure 12 (a). This plot suggests that *γ* = 1 is not a critical parameter for the timescale of the evolution either.

**Figure 12:**
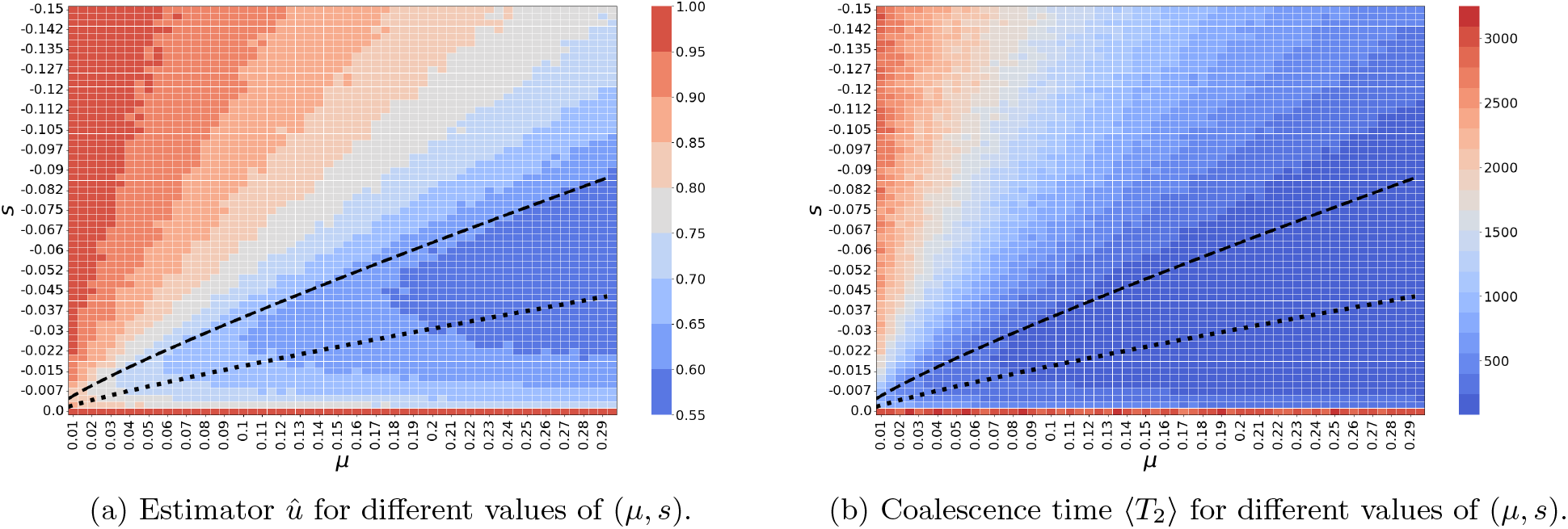
On both figures, the dashed line correspond to the line {*γ* = 1/2} and the dotted line to {*γ* = 1}.

#### Depth of the ancestry

In this section, we provide raw plots of the average pairwise coalescence time ⟨*T*_2_⟩. Figure 12 (b) highlights the drastic decrease in the timescale of evolution below the line {*γ* = ½} and suggests that it’s hard to distinguish polynomial and logarithmic scalings around the critical line.

#### The strong selection regime

In this section, we numerically compare the strong selection regime from [Cvi-jović et al., 2018] and the stable mutation-selection regime considered here. Let

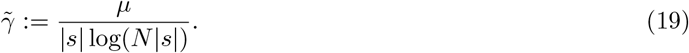

Then, note that the condition *N* |*s*| *e*^−*µ/*|*s*|^≫ 1 in [Cvijović et al., 2018] corresponds to the set 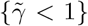 on the phase diagram; see Figure 13. On the other hand, we recall that the stable mutation-selection regime considered here is located above the {*γ* > 1/2} -line. It is thus clear from Figure 13 that this stable mutation-selection regime is a subset of the strong selection regime from [Cvijović et al., 2018].

**Figure 13:**
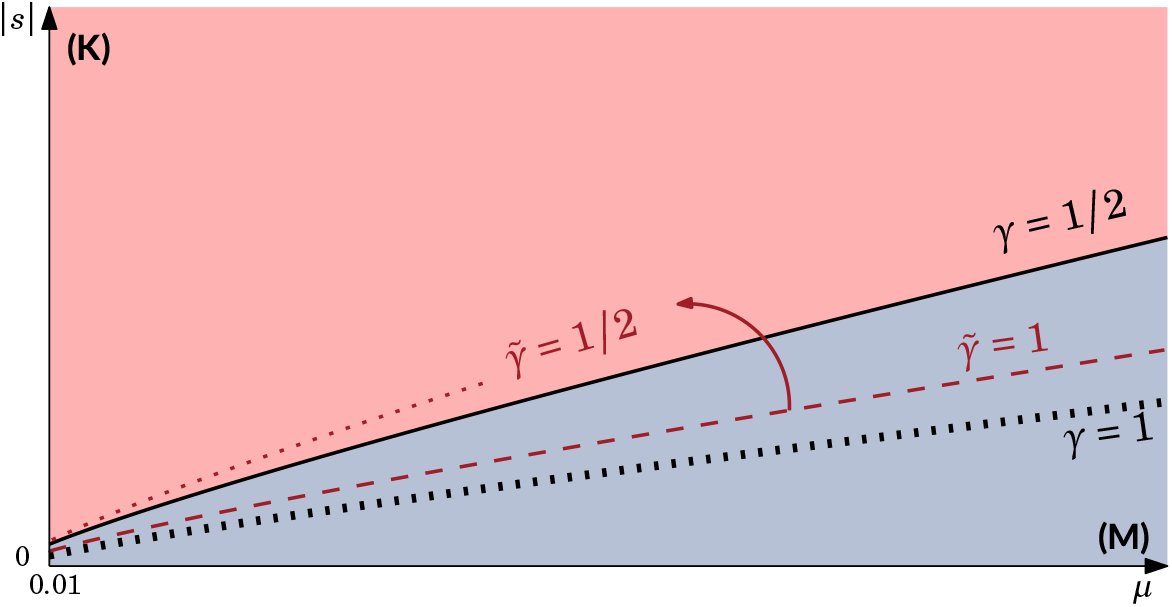
Phase diagram with level sets for the parameters *γ* (see (7)) and 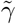 (see (19)).

### A.2 The distortion parameter

Recall from Section 4 that we write *T*_*k*_ for the time to the most recent common ancestor of a sample of size k and 𝔼 [*T*_*k*_] for the corresponding expectation. In this section, we derive the relation between the distortion parameter *α* and the expected coalescence times (𝔼 [*T*_*k*_]) given in (3).

#### Lemma A.1.

*Assume that the genealogical structure of the population is given by a Beta*(2 − *α, α*)*-coalescent. Then, we have*

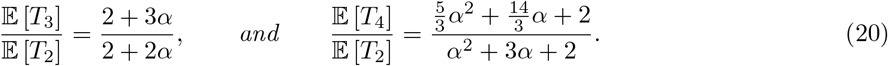

*In particular, we remark that for α* = 2 *(Kingman)*,

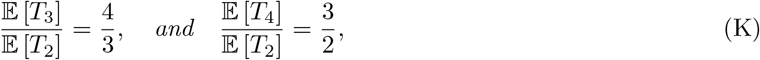

*for α* = 1 *(Bolthausen-Sznitman)*,

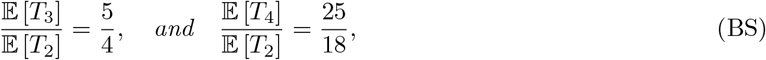

*and for α* = 0 *(Star-shaped)*,

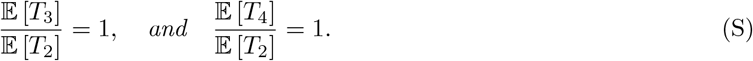

*Proof*. We give a general method that can easily be adapted to compute the ratios for *k* ≥ 5. Recall from (2) that the rate at which *k* blocks among *n* merge is given by

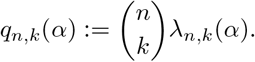

One can check (see e.g. [Gnedin and Yakubovich, 2007, Eq. (7)]) that the total rate at which a coalescence event occurs can be written as

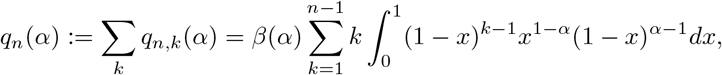

where *β* is as in (2). Let us now consider the Γ function defined by

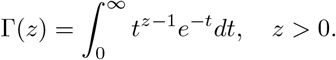

It is well-known that

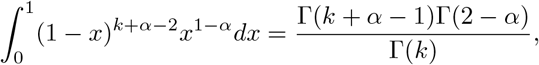

and that

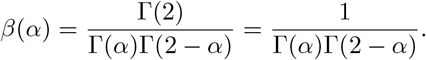

Putting all of this together and recalling that Γ(*z* + 1) = *z*Γ(*z*) for all *z* > 0, we get that

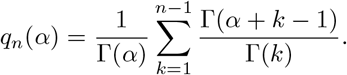

One can then check that (see e.g. [Birkner et al., 2023, Lemma A.1]) that the above equation reduces to

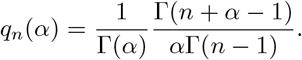

For *n* ∈ {2, 3, 4}, we thus get

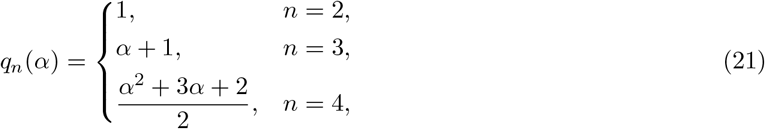

using the identity Γ(*z* + 1) = *z*Γ(*z*).

Let us now compute the expected coalescence times 𝔼 [*T*_*k*_]. Since *q*_2_(*α*) = 1, we have 𝔼 [*T*_2_] = 1. For *k* = 3, either all three individuals merge simultaneously, or there are two separate pairwise mergers. Let us denote by *P*_*n,i*_ the probability that, in a population of *n* individuals, the first coalescence event involves exactly *i* of them. Then, for *k* = 3, we have

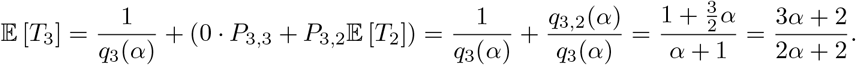

Similarly, we decompose *T*_4_ into the three possible outcomes for the first coalescence event and get that

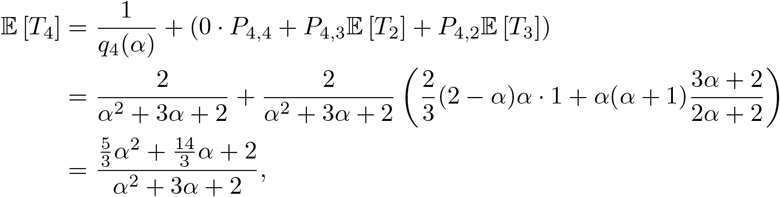

which concludes the proof of the lemma.

